# Neuronal actin dynamics, spine density and neuronal dendritic complexity are regulated by CAP2

**DOI:** 10.1101/027946

**Authors:** Atul Kumar, Lars Paeger, Kosmas Kosmas, Peter Kloppenburg, Angelika A. Noegel, Vivek Peche

**Author notes:** To whom correspondence should be addressed: Vivek S. Peche, Institute of Biochemistry I, Medical Faculty, University of Cologne, Joseph-Stelzmann-Str. 52, D-50931 Köln, Germany; Angelika A. Noegel, Institute of Biochemistry I, Medical Faculty, University of Cologne, Joseph-Stelzmann-Str. 52, D-50931 Köln, Germany.

## Abstract

Actin remodeling is indispensable for dendritic spine development, morphology and density which signify learning, memory and motor skills. CAP2 is a regulator of actin dynamics through sequestering G-actin and severing F-actin. In a mouse model, ablation of CAP2 leads to cardiovascular defects and delayed wound healing. This report investigates the role of CAP2 in the brain using *Cap2^gt/gt^* mice. Dendritic spine density and neuronal dendritic length were altered in *Cap2^gt/gt^*. This was accompanied by increased F-actin content and F-actin accumulation in cultured *Cap2^gt/gt^* neurons. In membrane depolarization assays, *Cap2^gt/gt^* synaptosomes exhibit an impaired F/G actin ratio, indicating altered actin dynamics. We show an interaction between CAP2 and n-cofilin, presumably mediated through the C-terminal domain of CAP2 and is cofilin ser3 phosphorylation dependent. In vivo, the consequences of this interaction were altered phosphorylated cofilin levels and formation of cofilin aggregates in the neurons. Thus, our studies identify a novel role of CAP2 in neuronal development and neuronal actin dynamics.

## Introduction

The coordinated regulation of assembly and disassembly of the neuronal actin cytoskeleton defines the range and complexity of morphologies in neurons. In particular, neurite outgrowth and polarity, the morphogenesis of dendrites and dendritic spines, synapse formation and stability require a functioning actin cytoskeleton ( Luo, 2002; Ahuja *et al.*, 2007; Rocca *et al.*, 2008). Axonal growth cone filopodia and dendritic filopodia, which are actin rich structures, play an essential role in synaptogenesis by directing and orchestrating pre and post synaptic factors at the synapse thereby acting as a functional bridge between neurons (Dailey and Smith, 1996; Evers et al., 2006; Fiala et al., 1998; Okabe et al., 2001). Subsequently, axonal growth cone filopodia develops to the pre synaptic terminal and the dendritic filopodia develops to post synaptic terminal, the dendritic spine. At the presynaptic terminal neurotransmitters are enclosed in synaptic vesicles which are docked at the presynaptic plasma membrane. These neurotransmitters are taken by neurotransmitter receptors on dendritic spine and postsynaptic density.

Actin binding proteins determine the G-/ F-actin ratio by regulating F-actin assembly and disassembly at the pre and post synaptic terminal. At the pre synaptic terminal actin orchestrates synaptic vesicle cycle leading to neurotransmitter release whereas at the post synaptic terminal it organizes and helps in trafficking post synaptic receptors and their scaffold. Pre and post terminal actin scaffold can be remodeled by altering actin dynamics (Fiala *et al.*, 1998; Cingolani and Goda, 2008; Hotulainen and Hoogenraad, 2010; Hotulainen *et al.*, 2009; Jontes and Smith, 2000). Furthermore, actin depolymerization by latrunculin A results in a near complete loss of synapses in hippocampal neurons (Zhang and Benson, 2001). It also affect growth cone development, neurotransmitter release, dendritic spine development and synaptic plasticity (Fukazawa et al., 2003; Sarmiere and Bamburg, 2004). Actin defines the spine morphology and spine density. Abnormalities in spine number and morphology were shown in many neurological disorders including intellectual disability, autism spectrum disorder and fragile X syndrome (Penzes et al., 2011; Van Spronsen and Hoogenraad, 2010). Taken together the actin cytoskeleton is an important component of a functional synapse and is tightly regulated to maintain the pool of F/G actin.

Cyclase associated proteins (CAP) are evolutionary conserved proteins which regulate the pool of actin monomers and also sever actin filaments. CAP1 shows a wide tissue distribution whereas CAP2 is primarily present in brain, heart and skeletal muscle, skin and testis. CAP1 has been reported as neuronal growth cone associated protein and as regulator of growth cone morphology through rearrangement of F-actin (Lu et al., 2011; Nozumi et al., 2009). Moreover, mammalian CAP1 was shown to be a proapoptotic protein (Wang et al., 2008). CAP2 sequesters G-actin and efficiently fragments filaments. This latter activity resides in its WH2 domain (Peche *et al.*, 2013). Earlier we reported on the function of CAP2 in the heart and in wound healing using mice lacking CAP2. During wound healing its loss leads to altered contractility. Moreover, ablation of CAP2 is lethal and mice lacking CAP2 develop a cardiomyopathy and have a disarrayed sarcomeric organization. Only a subset of mice survives and overcomes the lethal phenotype by an unknown mechanism (Kosmas *et al.*, 2015; Peche *et al.*, 2013). Cap2 is present at chromosome 6p22.3 in human, and an interstitial 6p22-24 deletion syndrome of the short arm of chromosome 6 was reported where patients with this deletion have a variable phenotype including a developmental delay, heart defects and cognitive dysfunction (Bremer et al., 2009; Davies et al., 1999). Given the abundance of CAP2 in the brain, its actin regulatory properties and restricted tissue distribution pattern, it may have a significant role in the brain and in particular for neuronal development.

Here we describe the function of CAP2 in the brain. CAP2 is expressed in different regions of the embryonic and adult brain. Using biochemical fractionation techniques, we found that CAP2 is a component of the presynaptic cytoskeletal matrix and crude synaptic vesicles. We show that CAP2 ablation leads to morphological differences in neurons with an increase in spine density and dendritic complexity. Furthermore, F-actin accumulated in the soma and neuritic processes. Membrane depolarization assays with isolated synaptosomes identified CAP2 as neuronal actin regulator. Interestigly, the level of phosphorylation of cofilin was decreased and was associated with its aberrant localization. Taken together, our data point to the importance of CAP2 in neuronal actin dynamics.

## Results

### CAP2 is expressed in the brain and localizes to dendrites and presynaptic terminal

For a detailed analysis of CAP2 expression in whole brain, we used the gene trap mice and followed the β-galactosidase fusion protein derived from the LacZ reporter and observed high expression in the olfactory bulb, cortex, hippocampus and cerebellum (Fig 1A). Western blot analysis with lysates from various brain regions at E18, P30 and P365 showed that the CAP2 levels were relatively low in the olfactory bulb and hippocampus at E18 whereas at P30 and P365 the levels were increased compared to E18 (Fig 1B). In contrast, CAP1 was present at relatively high levels in these parts of the brain at E18. However at P30 and P365 CAP1 was expressed uniformly in all regions of the brain (Fig 1C). Immunofluorescence analysis revealed CAP2 in the cortex, hippocampus and cerebellum (Fig. S1 A-C).

**Figure 1.**
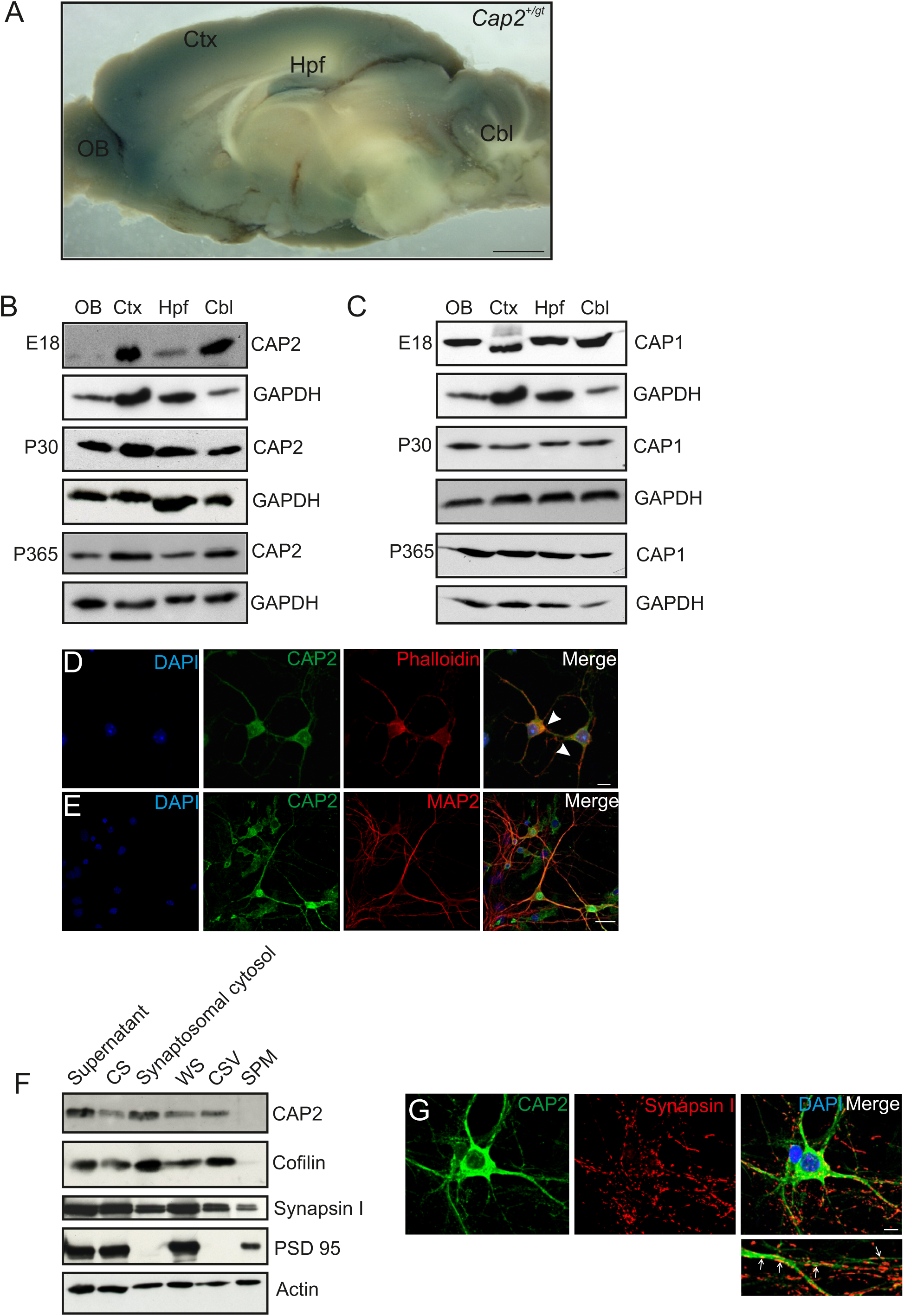
Expression and localization of CAP2 in brain. **(A)** X-gal staining for *Cap2^+/gt^* brain shows a widespread distribution of CAP2. The major anatomical regions of brain olfactory bulb (OB), cortex (Ctx), hippocampus (Hpf) and cerebellum (Cbl) are depicted. Scale bar, 1mm. **(B)** Western blot analysis of CAP2 in lysates from dissected brain regions at different developmental stage. **(C)** Western blot analysis of CAP1 in lysates from dissected brain regions at different developmental stage. **(D)** Immunocytochemistry of cultured cortical pyramidal neurons demonstrates the presence of CAP2 (green) in the cell body and in neurites. F-actin rich structures were visualized with TRITC-phalloidin (red). Scale bar, 10µm. **(E)** Co-labeling with Map2 (red), a marker of neuronal dendrites revealed the presence of CAP2 in dendritic shaft. Scale bar, 10µm. **(F)** CAP2 is present in synaptosome isolated form mice. CAP2 is located in the Synapsin I-positive synaptosomal cytosol, crude synaptic vesicle (CSV) region but not in the synaptic plasma membrane. Synapsin I (pre-synaptic marker) and PSD-95 (post synaptic marker) confirms the purity of the synaptosomal fraction. β-actin is used a loading control. **(G)** Co-labeling of cultured cortical neurons with CAP2 (green) and synapsin I (red) revealed the presence of CAP2 in presynaptic structure (white arrows, high magnification). Scale bar, 10µm.

To follow the neuronal localization of CAP2, we performed immunofluorescence analysis with cultured cortical pyramidal neurons which revealed a homogeneous distribution of CAP2 throughout the cytosol and in neurites. CAP2 was present in the proximity of F-actin in the neuronal cytoplasm (Fig 1D). Co-staining with MAP2 as a dendritic marker demonstrated the presence of CAP2 in dendrites (Fig 1E). Therefore, it is possible that CAP2 might influence the dendritic morphology, dendritic protrusions and spine development. Next, we employed a biochemical approach to study the subcellular localization of CAP2, which relies on differential centrifugation to separate proteins present at soluble extrasynaptic matrix, presynaptic matrix and at the pre-and postsynaptic plasma membrane. Western blot analysis revealed the presence of CAP2 in the crude synaptic vesicle and presynaptic matrix but not in the fraction containing the synaptic plasma membrane (Fig 1F). Synapsin I and PSD95 were used as a presynaptic marker and postsynaptic marker, respectively. Cofilin can be seen mainly in the soluble fraction as reported previously (Rust et al., 2010). Actin is present in all fractions and served as loading control. Immunofluorescence analysis of excitatory neurons with antibodies specific for synapsin I demonstrated the presence of CAP2 in synapsin I positive presynaptic terminals (Fig 1G). Given the importance of actin as the most prominent cytoskeletal protein at both the pre- and post-synaptic terminals and our data showing distribution of CAP2 in the presynaptic matrix, one can not rule out CAP2 as an important role in presynaptic processes.

### Altered spine density and dendritic complexity in CAP2 mutant neurons

CAP2 shows restricted tissue distribution and is also expressed in the brain. *Cap2^gt/gt^* mice develop cardiomyopathy, arrhythmia and die due to sudden cardiac arrest. Interestingly few mice overcome this phenotype but still developed mild cardiomyopathy (Peche *et al*., 2013). To our surprise, unlike cardiovascular system, no gross defects were found in *Cap2^gt/gt^* brain either at young or at old stages. The size of the brain remained unaltered (Fig 2A; S2A). Nissl staining on *Cap2^gt/gt^* and control brains revealed no gross differences in brain morphology (Fig 2B, S2B). Cortical layering, organization of hippocampus and the cerebellum architecture remained intact and unaltered (Fig 2C). CAP2 is present in the neuronal dendrites where actin is predominant and the degree of actin polymerization (F/G actin ratio) plays a crucial role in maintaining spine density and dendritic complexity (Hotulainen and Hoogenraad, 2010). To analyze the function of CAP2 in neuronal development, spine development and morphogenesis, we performed in vivo and in vitro analysis with WT and mutant neurons. We analyzed Purkinje neurons, which show high structural complexity when compared to other neuron types, and injected individual neurons in the cerebellum with the fluorescent dye biocytin at P60 and P365, followed by microscopic visualization. Confocal images revealed an increased dendritic complexity in mutant Purkinje neurons as compared to the WT (Fig 3A-D). Furthermore, we performed Sholl analysis on stacked confocal images using NeuronStudio for P60 (Fig 3E). Purkinje neurons from WT mice had a total dendritic length of ~4425 µm whereas CAP2 mutant mice showed an increase in dendritic length to ~5468 µm (Fig 3F; WT= 4425± 31 µm, n= 7; *Cap2^gt/gt^* = 5468± 335 µm, n= 9; P=0.04). The difference in dendritic length was more prominent in basal (WT= 768 ± 14 µm, n= 7; *Cap2^gt/gt^* = 1202± 85 µm, n= 9; P=0.005) and apo-basal regions (WT= 885± 90 µm, n= 7 neurons; *Cap2^gt/gt^* = 1077± 47 µm, n= 9 neurons; P=0.04) than in the apical region of Purkinje neurons. However, we did not observe a significant difference in the total dendritic surface area (Fig 3F; WT= 26910 ± 1617 µm^2^, n= 7; *Cap2^gt/gt^* = 29408 ± 2596 µm^2^, n= 9; P=0.385) and total dendritic volume (Fig 3F; WT= 17298 ± 148 µm^3^, n= 7; *Cap2^gt/gt^* = 13819 ± 2233µm^3^; n= 9; P=0.157) of CAP2 mutant Purkinje neurons.

**Figure 2.**
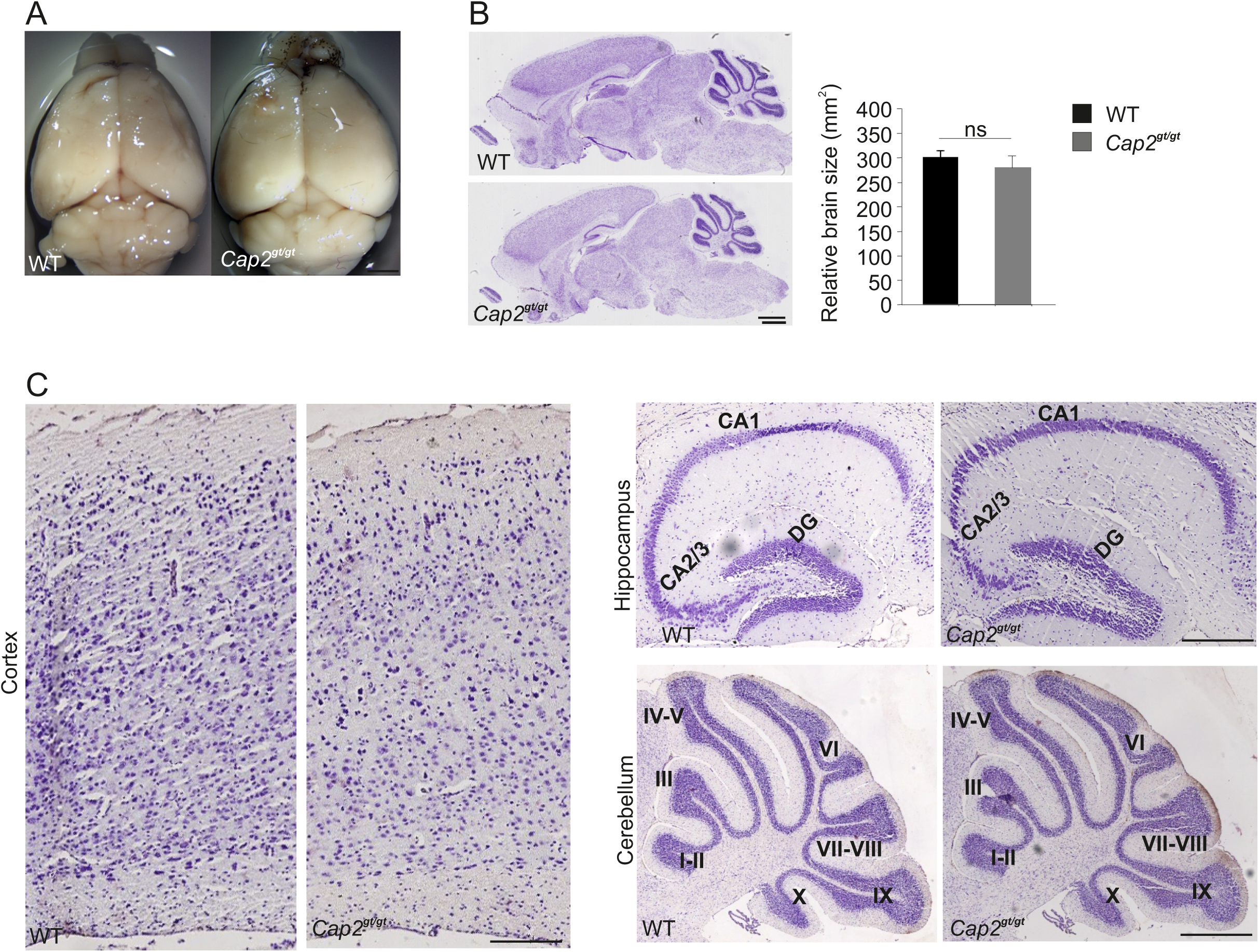
Morphology and Anatomy of *Cap2^gt/gt^* brain. **(A)** Comparison of brains isolated from wild-type mice as well as from homozygous does not reveal difference in the size of the brain (upper panel). Scale bar, 1mm. Cresyl violet staining of sagittal sections and its graphical analysis from control and mutant brains (middle panel) does not show any significant difference in gross anatomy of brain (WT= 300.68 ± 13.28 mm^²^ n= 10/3; *Cap2^gt/gt^* = 280.44 ±21.84 mm^²^ n=10/3 P = 0.3). Scale bar, 250µm. Higher magnification of indicated regions from cortical layering, hippocampus organization (cornu ammonis CA1-3; dentate gyrus DG) and foliation of cerebellum (folia I-X) is indicated (lower panel) Scale bar, 100 µm.

**Figure 3.**
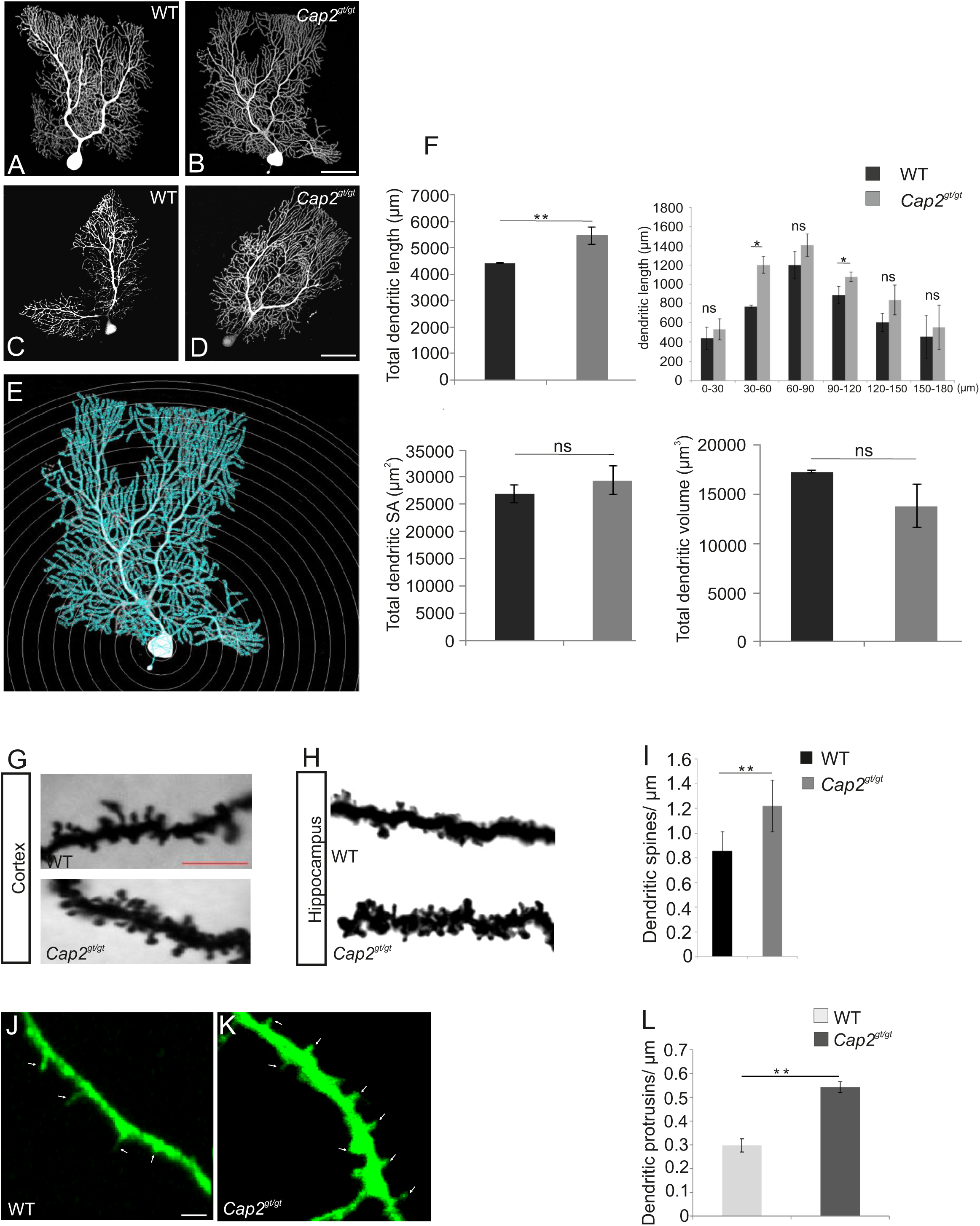
Dendritic spine density and neuronal complexity is altered in mutant neurons. **(A,B,C,D)** Purkinje neuron in cerebellum was injected with fluorescent dye biocytin at P60 (A,B) and P365 (C,D) to evaluate neuronal complexity and morphology in CAP2 mutant mice. **(E)** Sholl analysis using NeuronStudio was performed for P60. Scale bar, 20 µm. **(F)** Quantification of Sholl analysis for Purkinje neurons reveals an increase in total dendritic length but not in dendritic surface area and dendritic volume. **(G,H,I)** Golgi cox staining for WT and CAP2 mutant reveals an increase in spine density (WT= 0.85 ± 0.17 /µm; Cap2^gt/^*^gt^* = 1.2 ± 0.21 / µm). Scale bar: 20 µm. **(J,K,L)** Primary cortical neurons from WT and CAP2 mutant mice transfected with GFP show an increase number of dendritic filopodia in mutant neurons (WT= 0.3±0.05 spines/µm n=30/3; *Cap2^gt/gt^* =0.54 ± 0.036 n=24/3 P < 0.05).Scale bar: 5µm.

Golgi-Cox staining of the cortex and the hippocampus revealed a significant increase in spine density in *Cap2^gt/gt^* neurons (Fig. 3J-L; WT= 0.3±0.05 spines/µm; n=30 neurons/3; *Cap2^gt/gt^* =0.54 ± 0.036 spines/µm n=24 neurons/3 P < 0.01). Taken together, our data highlight the role of CAP2 in dendritic complexity, dendritic spine density and possibly also in synapse development.

### Actin filament turnover is impaired in CAP2 deficient neurons

The effects on dendritic complexity indicated CAP2 as an essential component in regulating actin dynamics during the maturation of neurons. To examine this possibility directly, we tested the F/G actin ratio in whole brain lysates from CAP2 mutant mice. We lysed forebrain tissue and performed fractionation assays to separate F-actin and G-actin. Western blot quantification revealed a 1.3 fold increase in F/G actin ratio in the CAP2 mutant brain lysate when compared to WT indicating an increase of F-actin in the mutant brain lysates (Fig 4A, WT= 1.06 ± 0.03, n= 4; *Cap2^gt/gt^* = 1.31 ± 0.08, n= 4; P << 0.05).

**Figure 4.**
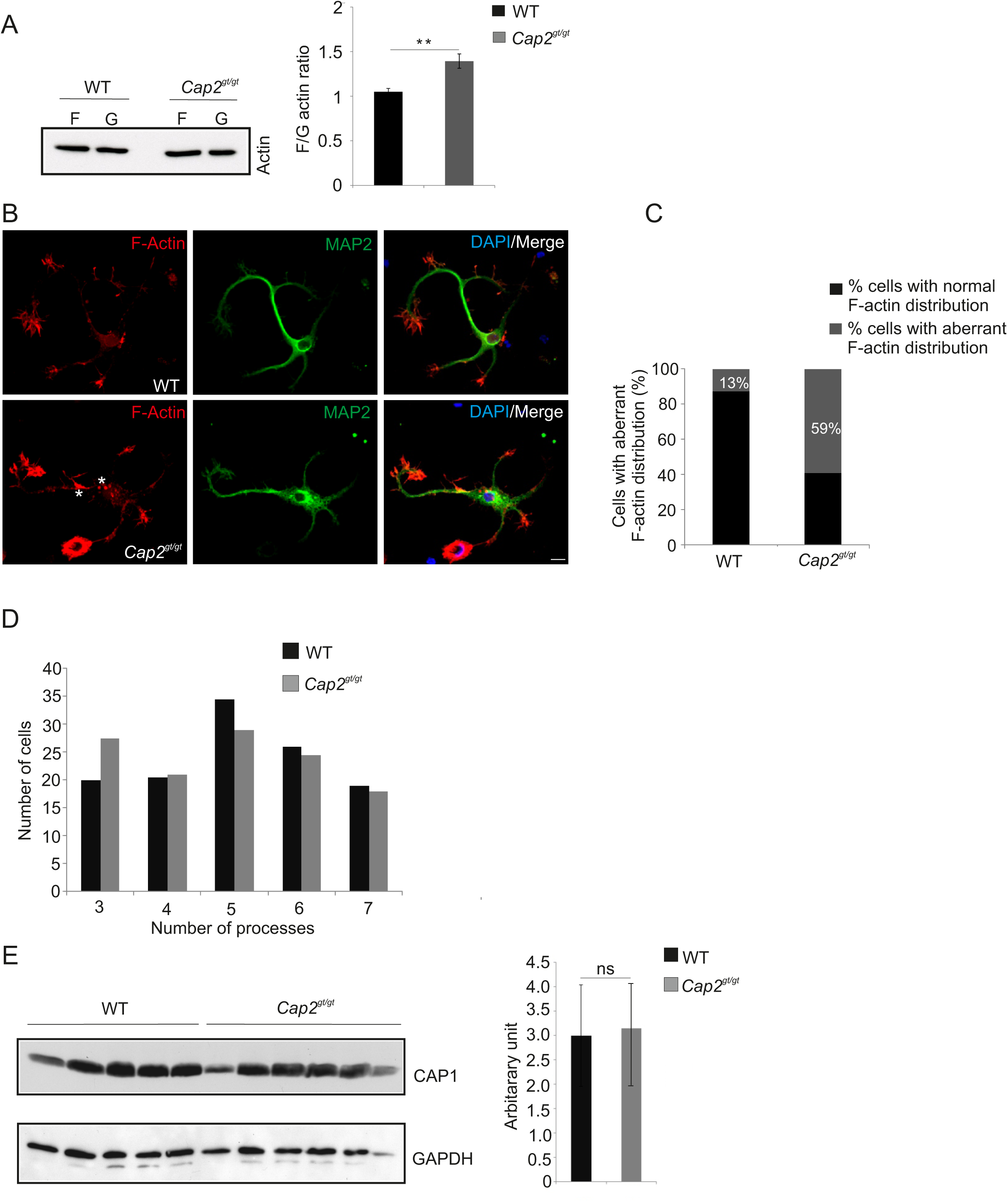
CAP2 deletion leads to actin accumulation and impaired actin dynamics. **(A)** Immunoblotting of WT and CAP2 mutant brain lysate for F/G-actin ratio reveals an increase in F-actin in mutant brain lysate (WT= 1.06 ± 0.03, n= 4; *Cap2^gt/gt^* = 1.31 ± 0.08, n= 4; P << 0.05). **(B)** Immunocytochemistry of cultured cortical neurons after 24 h in cultures demonstrated F-actin accumulation in soma and neuritic processes of mutant neurons. F-actin was visualized with phalloidin staining. Scale bar, 10µm. **(C)** Quantification of WT and mutant neurons showing aberrant F-actin distribution in the soma and/or neuritic processes. **(D)** Quantification of dendritic processes. The distribution of cells with a certain number of processes is shown. **(E)** Expression level of CAP1 in brain lysate was quantified after immunoblotting the WT and CAP2 mutant brain lysate with CAP1 polyclonal antibody (WT= 3 ±1.03 n= 5; *Cap2^gt/gt^* = 3.15 ± 1.05 n= 6; P>0.05).

In neurons, F-actin is primarily enriched at dendritic spines and forms a complex network which support the structures of dendritic spines (Cohen et al., 1985). Our initial observation of increased F/G actin ratio in mutant brain promted us to analyze the F-actin distribution microscopically in *in vitro* cultured neurons. In WT neurons, F-actin rich structures, detected with phalloidin staining, were predominantly present at the neurite growth cone. MAP2 was used as dendritic marker (Fig. 4B). Interestingly, unlike WT neurons, F-actin was accumulated in the soma as well as in the neuritic processes of mutant neurons (Fig. 4B, asterisk). We further counted cells with abnormal actin rich structures and found it to be statistically significant (Fig 4C, WT= 13% ± 3.7, n= 225 cells from 3 preparation; *Cap2^gt/gt^* = 59% ± 2.6, n= 225/3; P < 0.01). The alteration in the F-actin distribution prompted us to analyze neurite outgrowth. We could not observe a difference in the initial spreading of WT and mutant neurons within the first 24 h after plating as evident by the unaltered average number of processes per cell (Fig. 4D). Upon skin injury in CAP2 deficient mice, CAP1 was upregulated in the skin pointing towards a compensatory effect (Kosmas *et al*. 2015). To investigate a possible CAP1 compensatory effect in the brain, we checked the expression of CAP1 in WT and *Cap2^gt/gt^* mice brain lysates. Immunoblotting with CAP1 polyclonal antibodies did not reveal any difference in the brain CAP1 levels (Fig. 4E WT= 3 ±1.03, n= 5; *Cap2^gt/gt^* = 3.15 ± 1.05, n= 6).

### CAP2 regulates synaptic actin dynamics

Changes in actin dynamics have direct consequence on the efficacy of neurotransmitter release (Morales et al., 2000). To study the effect of CAP2 deletion on synaptic F-actin turnover, we performed fractionation studies with synaptosomes isolated from brain lysates of WT and CAP2 mutant cortex. An increase of 1.64 fold in the F/G-actin ratio was observed in CAP2 mutant synaptosomes when compared with the control (Fig 5A, WT= 1.1 ± 0.08, n= 4; *Cap2^gt/gt^* = 1.64 ± 0.13, n= 4; P << 0.05). The altered F/G actin ratio in mutant synaptosomes prompted us to investigate the synaptic actin dynamics. F-actin assembly and disassembly is a characteristic feature of depolarization of synaptosomes. With the stimulus, the F/G actin ratio undergo a rapid increase (Bernstein and Bamburg, 1989; Yamada et al., 2009). To investigate whether CAP2 has an effect on stimulated actin dynamics, we isolated forebrain synaptosomes and depolarized them with 40 mM KCl. This stimulus induced a rapid cycling of actin in control preparation as observed by a rapid assembly of actin at 3 seconds after the stimulus followed by a rapid disassembly at 10 seconds (Fig 5B). In accordance with earlier reports, the F/G ratio was increased with time when followed till 60 seconds (0 s: 1.09 ± 0.06, 60 s: 1.61 ± 0.14; n = 4 preparations from 4 different mice) (Bernstein and Bamburg, 1989; Wolf et al., 2015; Yamada et al., 2009). In mutant preparations, we observed an increased basal value of F/G actin ratio and no rapid actin assembly and disassembly at all-time points examined after the stimulus (0 s: 1.90 ± 0.20, 60 s: 2.28± 0.25, n = 4 preparations from 4 different mice). Notably after the stimulus, the F/G actin ratio was consistently higher than its corresponding control values (Fig 5B). When we quantified the values at time point 0 s till 60 s, we found an ~19 % increase in the F/G actin ratio which is significantly lower when compared to the control in which it was increased by ~47%. As reported previously (Bernstein and Bamburg, 1989) this controlled assembly/disassembly of actin is crucial for vesicle release. Our data indicates an aberrant actin dynamics upon a stimulus in mutant synaptosomes, which could have an effect on vesicle release. Taken together, our data highlights a role for CAP2 in synaptic actin remodeling.

**Figure 5.**
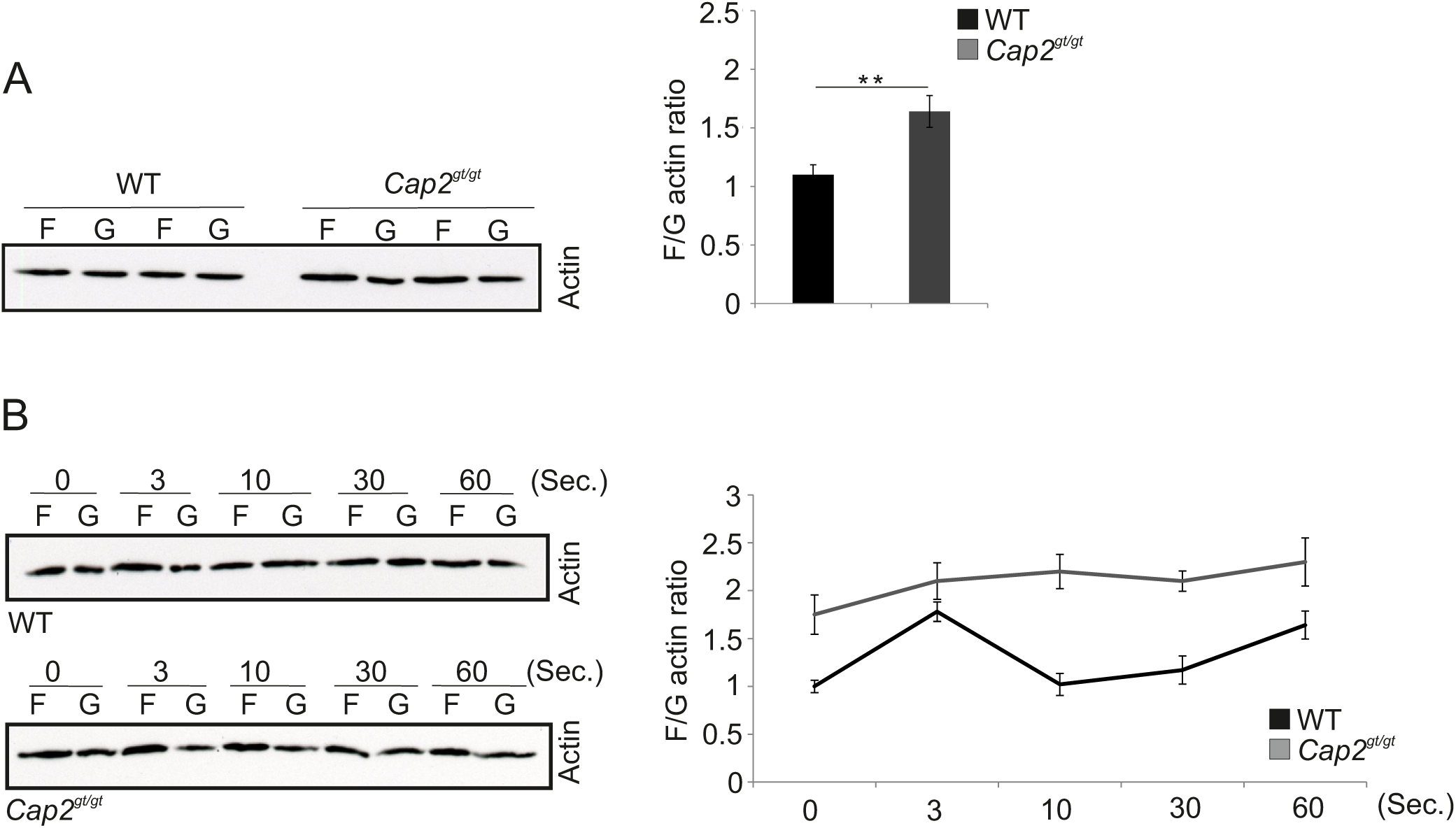
Synaptic actin dynamics is impaired in CAP2 null neurons. **(A)** Immunoblot showing the F-actin and G-actin fraction from a control and CAP2 mutant mice forebrain synaptosome. CAP2 mutant synaptosome shows an increase in the F/G-actin ratio when compared with the WT synaptosome (WT= 1.1 ± 0.08, n= 4; *Cap2^gt/gt^* = 1.64 ± 0.13, n= 4; P << 0.05). **(B)** Immunoblot showing the F-actin and G-actin fraction after KCl depolarization assay in a time dependent manner to WT and *Cap2^gt/gt^* forebrain synaptosome reveals a ~19 % increase in F/G actin ratio (0 s: 1.91 ± 0.20, 60 s: 2.28 ± 0.25 n = 4/4) which is drastically reduced when compared to the control in which it was increased by ~47% (0 s: 1.09 ± 0.12, 60 s: 1.61 ± 0.14 n = 4/4).

### CAP2 interacts with n-cofilin

Previous studies reported that CAP accelerates the dissociation of ADP G-actin–cofilin complexes, recharges actin monomers with ATP and releases cofilin from ATP G-actin complex for actin polymerization (Moriyama and Yahara, 2002). CAP1 interacts with cofilin in muscle and non-muscle cells. The interaction between CAP2 and cofilin has not been addressed so far. We performed supernatant depletion pull down assays with variable concentration of bead-immobilized GST-CAP2 and purified n-cofilin. A decrease of cofilin in the supernatant was observed which shows an interaction between CAP2 and n-cofilin (Fig 6A). The assay was carried out at physiological pH and salt concentration (Fig 6B). Next we narrowed down the interacting domain of CAP2 by performing pull down experiments with ectopically expressed GFP-cofilin and GST-tagged CAP2 polypeptides (Peche *et al.*, 2013). Pull-down assays revealed an interaction of full length CAP2 with n-cofilin while N terminal and WH2 domain did not show any interaction; a weak interaction was observed with a truncated protein lacking the first 55 amino acids of CAP2 and of the C terminal domain of CAP2 with n-cofilin. The WH2 domain did not substantially affect the C-CAP2/n-cofilin interaction (Fig 6C). This point at the importance of the C terminal domain of CAP2 for the interaction with n-cofilin. The CAP2-n-cofilin interaction was also supported by colocalization studies with WT primary cortical neurons where we found partial colocalization of n-cofilin and CAP2 at the cell periphery as well as in the cytoplasm and neurites (Fig 6D).

**Figure 6.**
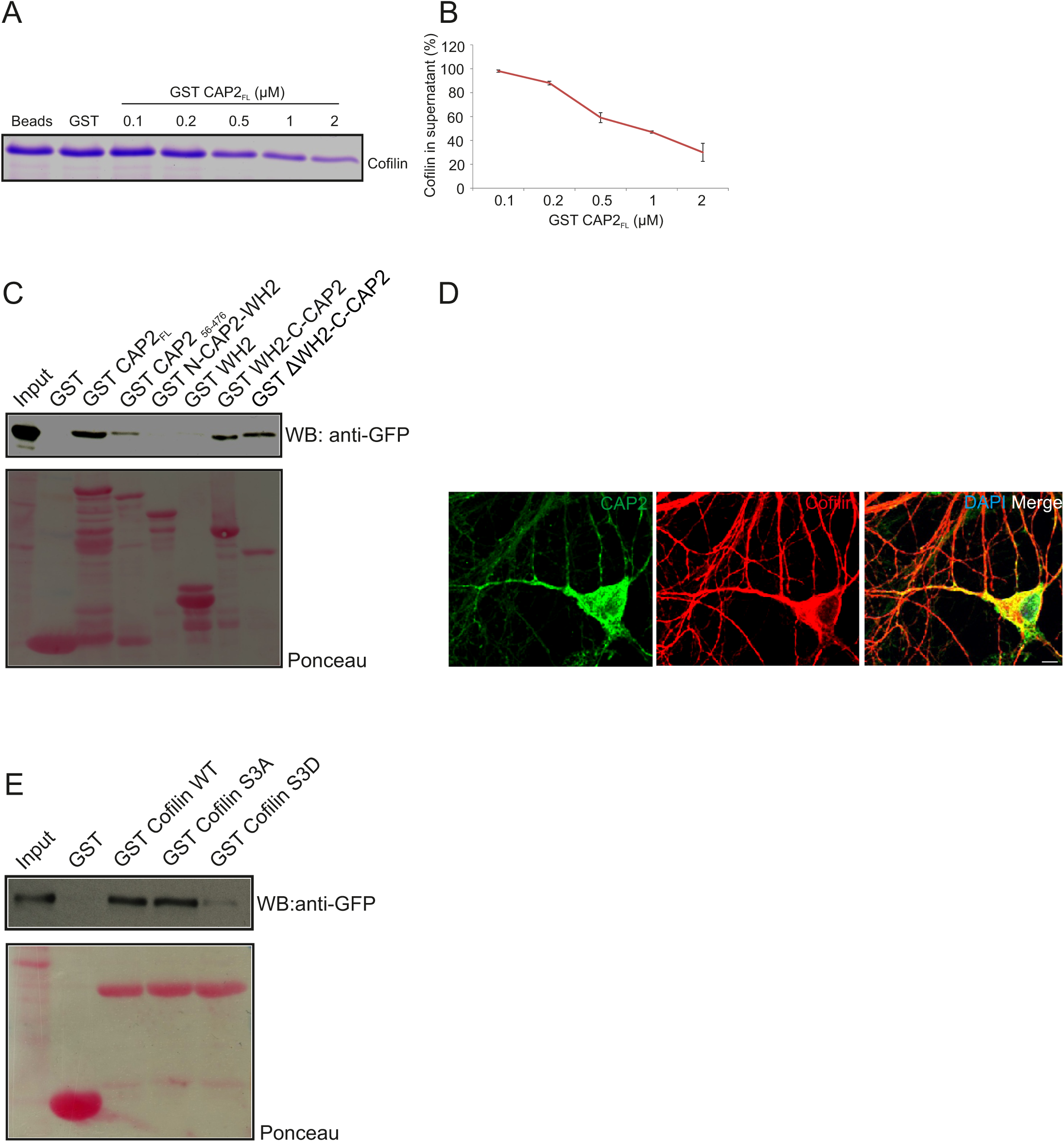
CAP2 pull-down n-cofilin through its C-terminal domain containing WH2. **(A)** Coomassie-stained SDS-polyacrylamide gel shows the supernatant depletion pull down assay for examining the interaction between the GST-CAP2 and thrombin cleaved cofilin protein from GST-cofilin protein. **(B)** The amount of cofilin in supernatant fractions was quantified from Coomassie-stained SDS-polyacrylamide gels by densitometry analysis using Image J (GST = 1; 0.1 µM = 0.98±0.01; 0.2 µM = 0.88±0.015; 0.5 µM = 0.59±0.04; 1 µM = 0.47±0.01; 2 µM = 0.3±0.07; n=3). **(C)** Pull-down experiment with ectopically expressed GFP N-cofilin. Precipitated proteins were detected by western blotting using monoclonal anti-GFP. Ponceau staining as loading control of nitrocellulose membrane for pull down assay. GST was used as a control. CAP2 interacts with N-cofilin through its C terminal which includes WH2 domain. **(D)** Immunocytochemical analysis by co-labeling with monoclonal CAP2 (green) and polyclonal cofilin I (red) revealed the colocalization in adult brain section. **(E)** Immunoblot of pull down using bacteria purified GST-n-cofilin WT, GST-cofilin S3A mutant showed a pull down of CAP2 from the lysates whereas GST-n-cofilin S3D which mimics a phosphorylated Ser3 residue had reduced binding efficiency to CAP2. Ponceau staining as loading control of nitrocellulose membrane for pull down assay. GST was used as a control.

Cofilin exists in a phosphorylated and an unphosphorylated state, the latter being the active state (Sarmiere and Bamburg, 2004). Hence to analyze whether CAP2 binds to the phosphorylated (inactive) or unphosphorylated (active) n-cofilin, we used mutant proteins carrying mutations of serine 3 mimicking the phosphorylated (S3D) or unphosphorylated state (S3A) of cofilin. GST-WT n-cofilin and GST-n-cofilin S3A mutant pulled down CAP2 from lysates of GFP-CAP2 expressing HEK293T cells, whereas GST-n-cofilin S3D showed very weak binding to CAP2 (Fig 6E). Thus CAP2 has a potential to interact with n-cofilin and the activation states of cofilin also have an effect on this interaction.

### CAP2 affects the phosphorylated n-cofilin levels and leads to its aggregation

The CAP2 - n-cofilin interaction directed us to examine the effect of this interaction on cofilin regulation in neurons. The levels of phosphorylated (Ser3) n-cofilin were lower in whole brain lysates of *Cap2^gt/gt^* mice compared to WT (Fig 7A, B). Supporting this observation, immunofluorescence analysis revealed low levels of phospho (Ser3) cofilin in the brain. The staining with cofilin phospho-specific antibody showed reduced staining in the mutant cortex compared to WT (Fig 7C, D). The total n-cofilin was unaltered. We next examined a possible alteration in total n-cofilin localization in CAP2 mutant brain. In WT, n-cofilin was diffusively expressed in the cytosol (Fig 7E) whereas in mutant brain, we found n-cofilin accumulation in cytoplasmic aggregates, where it is presumably inactive (Fig 7F, asterisk). We also analyzed the expression of kinases and phosphatases involved in cofilin phosphorylation and dephosphorylation. Western blotting of phospho LIMK1/LIMK2 and LIMK2 did not show significant differences in their amounts in WT and CAP2 mutant brain lysates. The expression of TESK1 and ROCK1, kinases involved in n-cofilin phosphorylation, were also unaltered. Moreover, the levels of phosphatase chronophin remained unchanged (Fig S3). Taken together these results confirm that ablation of CAP2 leads to increased levels of dephosphorylated cofilin, along with n-cofilin aggregation.

**Figure 7.**
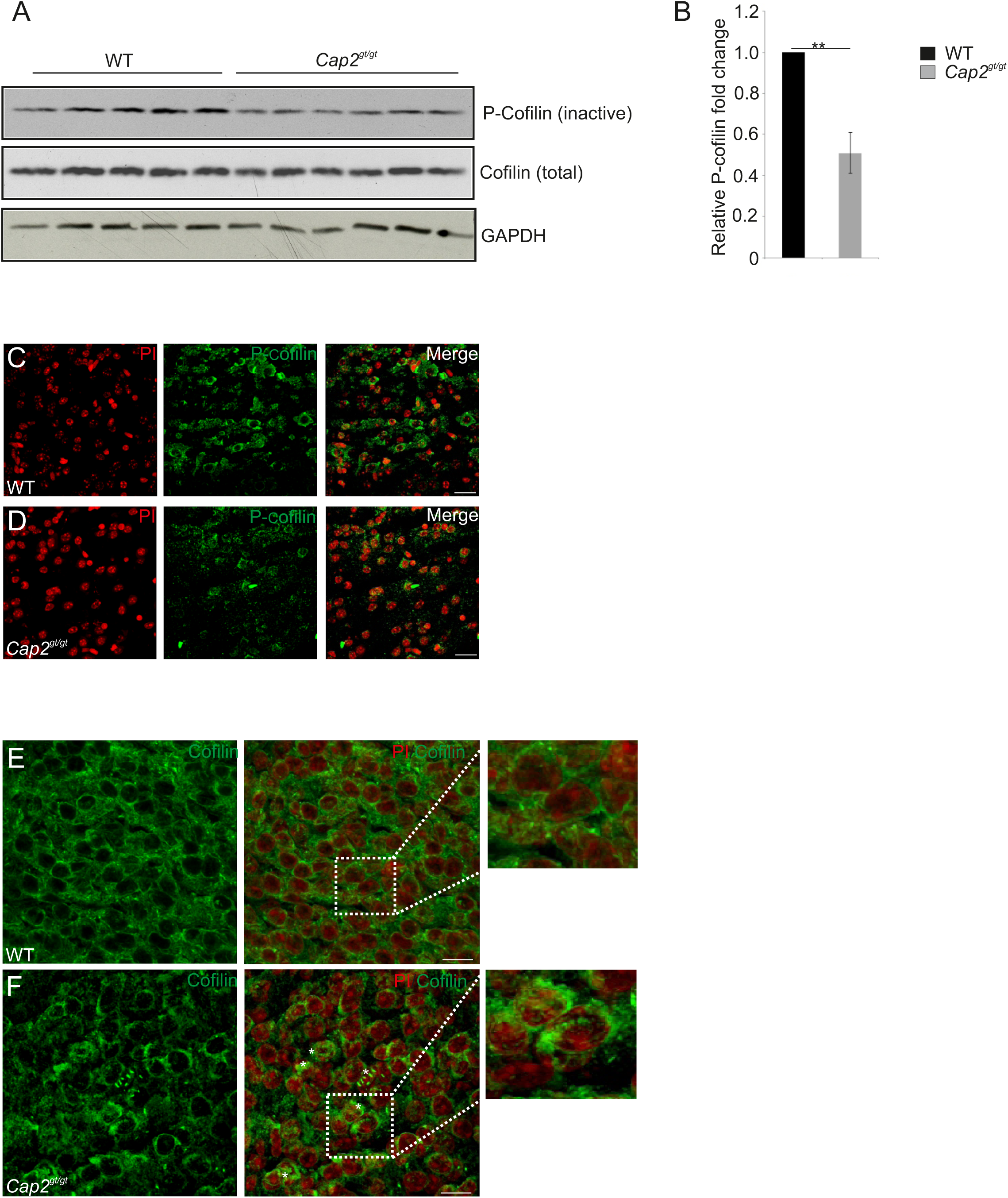
CAP2 depletion increase cofilin dephosphorylation and its accumulation. **(A)** Western blotting of phospho Ser 3 cofilin shows reduced cofilin phosphorylation in the CAP2 mutant brain lysate. Total cofilin expression was unchanged in WT and mutant brain lysate. GAPDH served as a loading control. **(B)** Quantification of cofilin phosphorylation from 6 independent experiments analyzed using Student’s *t*-test (WT = 1; *Cap2^gt/gt^* = 0.51 ± 0.1; n=6; P= 0.03). Values were normalized to loading control. **(C,D)** Immunohistochemical analysis of WT and CAP2 mutant brain section with rabbit monoclonal phospho cofilin antibody reveal a decrease in staining in cortical region of mutant brain. Scale bar: 10µM. **(E,F)** WT and Cofilin brain section was stained with polyclonal cofilin antibody. Accumulation of cofilin as cytoplasmic aggregates was observed in CAP2 mutant brain as indicated by arrow heads, whereas cofilin was nicely distributed in the cytoplasm and nucleus of WT brain. Scale bar: 10µM.

## Discussion

The actin cytoskeleton plays a major role in the morphology and structural changes of neurons. The equilibrium between G and F – actin regulates many cellular processes and the pool of G and F – actin is maintained by many actin binding proteins. CAP2 is a G-actin sequestering protein and also shows a F-actin severing potential, making it an important regulator of actin dynamics (Peche et al., 2007; Peche et al., 2013a). We reported previously that CAP2 knockout mice develop dilated cardiomyopathy (Peche *et al.*, 2013) and it plays a role in skin repair during wound healing, whereas its ablation leads to changes in infiltration of inflammatory cells and contraction of wounds (Kosmas et al., 2015).

In mature neurons, actin is the most prominent cytoskeletal protein at both the pre- and postsynaptic terminals. From its subcellular organization, actin has been implicated in maintaining and regulating vesicle pools (Cingolani and Goda, 2008). Upon biochemical fractionation, we observed the association of CAP2 with the pre synaptic matrix and crude synaptic vesicle. Although many reports point at an importance of actin regulation at the synapse, very few proteins were shown to directly affect the synaptic actin dynamics making it an elusive mechanistic event. Here we provide evidence that ablation of CAP2 results in increased synaptic F-actin levels and regulates synaptic actin polymerization. Moreover, we show that it is involved in regulating actin dynamics in isolated synaptosome. Previously ADF, n-Cofilin and Profilin2 have been reported to regulate synaptic actin dynamics and neurotransmitter release. For n-cofilin knockout mice increased spine density was reported, whereas the ADF/n-cofilin double knockouts showed decreased spine density. Moreover, in the double mutant synaptic actin dynamics was impaired and more severely affected than in the single mutants. Interestingly, genetic ablation of ADF had no adverse effects on synapse structure or density which was explained by elevated n-cofilin levels in ADF mutants (Pilo Boyl *et al.*, 2007; Görlich *et al.*, 2011; Wolf *et al.*, 2015). Our data revealed a presynaptic role for CAP2 suggesting it to be a further controlling factor in neurotransmitter release and neuronal excitability.

An increase in F-actin was observed in lysates from CAP2 mutant brain. F-actin staining in cultured WT neurons showed F-actin rich structures predominantly at the neuritic growth cone, whereas actin punctae were observed in the soma and neuritic processes of mutant neurons. Depletion of CAP1 also leads to a heavy accumulation of abnormal F-actin structures in NIH/3T3, N2A and HeLa cells (Bertling et al., 2004; Zhang et al., 2013). F-actin aggresomes have been reported in neuronal cells which comprise an aberrant accumulation of multiple fragments of F-actin trapped in the cytoplasm (Lázaro-Diéguez et al., 2008). Similar punctae were also observed in CAP2 mutant neurons, which points at the role of CAP2 in maintaining cellular F/G actin ratios. The imbalance in this ratio could ultimately lead to the formation of these structures, which are detrimental to the cell depending on their composition and abundance.

Neuronal dendrites develop a small-specialized actin rich structure known as dendritic spine. At the structural level, more stable dendritic spines are developed from the highly motile dendritic filopodia-like structures. Actin nucleation by Arp2/3 and the formin mDia2 together with actin filament disassembly by ADF/cofilin promote localized actin filament turnover. This generates a mechanical force against the dendritic plasma membrane that can lead to initiation and elongation of dendritic filopodia (Honkura et al., 2008; Hotulainen et al., 2009). In CAP2 mutant neurons we observed a significant increase in dendritic filopodia formation at div 7. Furthermore, Golgi Cox staining in WT and *Cap2^gt/gt^* adult brain revealed a significant increase in dendritic spines in mutants. Similar observations were made when n-cofilin was deleted from principal neurons of the postnatal forebrain, which resulted in an increased synaptic F-actin content, increased spine density and enlargement of dendritic spines (Rust et al., 2010). In various studies it has been shown that dendritic spines are also eliminated by spine pruning. From our observation of CAP2 in neuronal dendrites and CAP2 being a regulator of actin filament dynamics, a role of CAP2 in dendritic filopodia formation and maintenance of dendritic spines is likely. Additionally, in CAP2 mutant Purkinje neurons we observed a difference in dendritic complexity. Although we did not observe a compensatory effect of CAP1 upon CAP2 deletion, this does not rule the possibility of alternative mechanisms even by unchanged levels of CAP1 by which the severity of the Purkinje neuron phenotype is attenuated. Similar observations were also made in the protein family of phosphatidylinositol 3,4,5- triphosphate dependent Rac exchanger 1 (P-Rex1) and P-Rex2. A double knock out of P-Rex1 and P-Rex2 showed a very strong Purkinje cell dendritic morphology phenotype and motor neuron coordination defects in comparison to the mild phenotype of the P-Rex2 knockout (Donald et al., 2008). Very little is known about the molecular mechanism behind the interstitial branching in dendrites. However it is thought that destabilization of cortical actin and the membrane can lead to filopodial protrusions, which act as precursors of transient branches.

Cyclase associated protein actively cooperates with cofilin and actin to regulate actin dynamics (Hubberstey and Mottillo, 2002; Moriyama and Yahara, 2002; Ono, 2013; Zhang et al., 2013). The nematode *C*. *elegans* also has two CAP isoforms; CAS-1 and CAS-2 where CAS-1 binds to actin monomers and enhances exchange of actin-bound ATP/ADP even in the presence of UNC-60B, a muscle-specific ADF/cofilin; CAS-2 strongly enhances the exchange of actin bound nucleotide in the presence of UNC-60A, a *C*.*elegans* non muscle-specific ADF/cofilin (Nomura and Ono, 2013; Nomura et al., 2012). Our data demonstrates that CAP2 also interacts with n-cofilin. Moreover this interaction, unlike CAP1, resides in the C-terminal domain of CAP2. CAP1 and CAP2 primarily differ in their N-terminal domain, which could be one possible explanation for CAP2’s C-terminal interaction with n-cofilin. Interestignly, CAS-2 N-terminal does not interact with G-actin-UNC-60A complex suggesting it to behave similarly as mouse CAP2 (Nomura and Ono, 2013). We also report that ablation of CAP2 in neuron results in the reduction of phosphorylation of cofilin and its accumulation in cytoplasmic aggregates. However reduced expression of various kinases like LIMK, TESK or ROCK was not observed in CAP2 mutant neurons. This is in agreement with earlier reports where CAP1, the other isoform of CAP, when knocked down gave rise to reduced cofilin phosphorylation and cofilin aggregation (Bertling et al., 2004; Zhang et al., 2013).

Many psychiatric and neurological disorders like Autism spectrum disorder (ASD), Schizophrenia and Alzheimer’s are accompanied by alterations in spine number and dendritic complexity (Penzes et al., 2011; Shirao and González-Billault, 2013). Altered neuronal actin dynamics lead to behavioral phenotype in mice (Rust and Maritzen, 2015). *Cap2^gt/gt^* mice showed a reduced performance during the rotarod test and significantly decreased rearing activity. Interestingly, CAP2 is important for motor coordination but not for motor learning as performance of Cap2^gt/gt^ mice during rotarod tests improved after subsequent trials but latency to fall was higher in mutant mice. In independent studies conducted at the international mouse phenotyping consortium (IMPC), Cap2 heterozygous animals also show decreased rearing, implying a role of CAP2 in altering the ability to initiate locomotor activity (p=0.0038). Furthermore, rota rod analysis of Cap2 heterozygous animals at the IMPC also revealed impaired proprioception measured through rotarod tests, but in contrast to homozygous animals, heterozygous animals showed reduced latency to fall as compared to WT (p=0.02). Further investigations with a brain specific CAP2 knockout must be done in order to conclude the brain specific function of CAP2 in the behavioral phenotypes observed.

n-cofilin deletion from principal neurons of the postnatal forebrain impairs synaptic plasticity during associative learning (Rust et al., 2010). Additionally, double mutant mice lacking ADF and n-cofilin, ADF-/-/n-Cof^flx/flx,CamKII-cre^, were shown to develop attention-deficit hyperactivity disorder (ADHD). The proposed hypothesis for the development of ADHD-like phenotype in these mice involves impaired actin dynamics resulting in altered neurotransmitter release (Zimmermann et al., 2015).

CAP2 is a gene of interest for possibly contributing to both the cognitive delay and the heart defects in patients with a chromosome 6p22 deletion (Bremer et al., 2009). Further, a number of genes involved in dopamine receptor mediated signaling pathways were dysregulated in schizophrenics. These include genes in the protein kinase A pathway (protein kinase ARII subunit, adenylyl cyclase-associated protein 2) and protein kinase C (Hakak et al., 2001). Also, upregulation of CAP1 has been reported in the brain of schizophrenic patients (Martinsde-Souza et al., 2010). This points towards a role of CAPs in neurological disorders. Therefore it will be interesting to decipher the role of CAP2 in various human neurological disorders. Further studies with human samples are needed to find possible mutations in CAP2 to support these findings.

In conclusion, depletion of CAP2 leads to an increase of F-actin in CAP2 mutant neurons along with an increase in dephosphorylation of cofilin and the appearance of cytoplasmic cofilin aggregates. The interaction of CAP2 and n-cofilin could be essential to keep n-cofilin in its unaggregated state thereby protecting neurons from pathological consequences. Increased F-actin levels in mutant neurons is the most probable cause of the morphological defects observed which include increased dendritic spine density and neuronal complexity (Fig 8). Altered actin dynamics in CAP2 deficient synaptosomes could also lead to an impaired neurotransmitter release, which must be addressed further. Thus, CAP2 is an important actin regulator in the brain and its deletion leads to several pathological consequences that are also found in neurological disorders.

**Figure 8:**
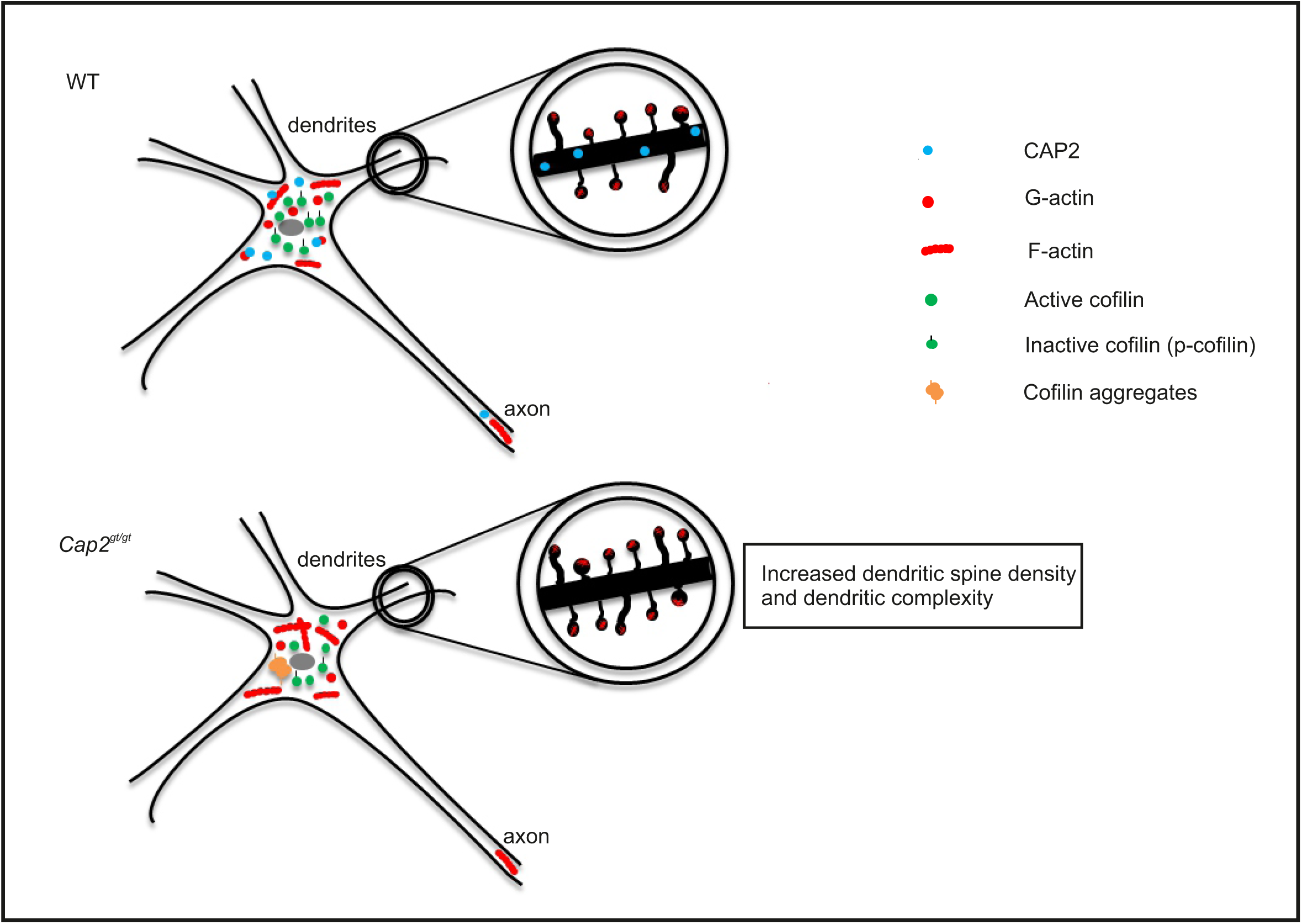
Model illustrating the role of CAP2 in neurons. CAP2 sequesters G-actin and severs F-actin to regulate actin dynamics. When it is absent, F-actin is increased. Due to the increase in F-actin and possibly through the loss of CAP2-cofilin interaction in mutant neurons, dendritic spine density and dendritic complexity is altered. Another consequence of the loss of CAP2 is the formation of cofilin aggregates in the cytoplasm of neurons which could be pathological.

## Materials and methods

### Transgenic animals

CAP2 deficient mice generated by a gene trap approach have been described previously (Peche et al., 2013). C57/Bl6 wild type (WT) mice derived from the breeding of *Cap2^gt/+^* animals were used as controls.

The animals had free access to water and standard laboratory chow diet and were kept at an artificial light/dark cycle at 20-22°C. All procedures were performed in accordance with the animal protection law stated in the German civil code and were conform to the Guide for the Care and Use of Laboratory Animals published by the US National Institutes of Health (NIH Publication No. 85-23, revised 1985).

### Histology

Embryonic (E18.5) and juvenile mice (P8-30) were perfused with 4% paraformaldehyde (PFA). Brains were fixed in the same fixative for 4-6 h. Next day the brains were processed for paraffin embedding. Sections were cut at 10 µm using a microtome, collected in deionized water and transferred onto slides.

### Immunohistochemistry, antibodies and histology

For general histology, the samples (paraffin sections) were stained with creysl violet according to standard procedures. Images were acquired on a Leica SCN400 slidescanner with autoloader. LacZ staining for the *Cap2^+/gt^* brain was done as described previously (Peche et al., 2013). For immunofluorescence, paraffin sections were deparaffinised in 2 changes of xylene and rehydrated through a graded ethanol series, which was then followed by antigen retrieval and antibody incubation. For brain samples incubation was done with monoclonal anti-CAP2 (1:50) (Kosmas et al., 2015), polyclonal anti-cofilin (1:50; cell signaling), polyclonal anti-phospho cofilin (1:50, cell signaling), monoclonal anti-β-actin (1:25; clone AC 74, Sigma) antibodies. Appropriate secondary antibodies conjugated with Alexa Fluor 488 and 568 (Molecular Probes) were used. Nuclei were visualized with either 40,60-Diamidino-20-phenylindole (DAPI) or propidium iodide (PI). Sections were mounted and imaged with a Leica SP5 confocal microscope.

### Isolation of primary cortical neurons

Primary mouse cortical neurons were dissected from E17.5-E18 brains and cultured as previously described (Beaudoin et al., 2012). The cortical region of the brain was dissected in a dissection medium containing HBSS (97.3%, Ca^2+^ and Mg^2+^ free), sodium pyruvate (1x), glucose (0.1% w/v), HEPES (pH 7.3, 10 mM), trypsinized and dissociated individually for each embryo to maintain separate genetic identities of the neuronal cultures. Neurons were plated at densities between 100-150 cells/mm^2^ on poly L-lysine coated glass coverslips and cultured at 37°C and 5% CO_2_ in Neurobasal medium 96% (Invitrogen) supplemented with B27 (1%, Invitrogen), 200 mM glutamine(1%) and Penicillin/streptomycin (1%). Cultured cells were fixed with paraformaldehyde and processed for immunofluorescence analysis using anti-CAP2 polyclonal (1:100) (Peche et al., 2007), anti-CAP2 monoclonal, anti-MAP2 polyclonal (1:50; NEB), anti-synapsin 1 polyclonal (1:200; Thermo Scientific), anti-PSD-95 monoclonal (7E3-1B8, 1:200; Thermo Scientific) antibodies. F-actin was visualized with TRITC Phalloidin. Nuclei were stained with DAPI. Cells were incubated with primary and secondary antibodies for 1 h at room temperature each. Neurons were transfected with plasmids carrying cDNAs fused to GFP sequences using the Amaxa Neuron transfection kit (Lonza) according to the manufacturer’s specification for in vitro dendritic filopodia analysis experiment. Neurons were mounted and imaged with a Leica SP5 confocal microscope.

### Synaptosomal preparation

Synaptosomes from forebrain tissue from P30 mice were isolated as previously described (Blackstone et al., 1992). Briefly, tissue was homogenized in 10 volumes of HEPES-buffered sucrose (0.32 M sucrose, 4 mM HEPES, pH 7.4) with a glass Teflon homogenizer. Homogenates were centrifuged at 1000x g, 4°C, to remove the nuclear fraction. The resulting supernatant was further centrifuged at 10,000x g for 15 min to yield the crude synaptosomal pellet. The supernatant containing the cytosolic and light membrane fraction was collected. The pellet was resuspended in 10 volumes of HEPES-buffered sucrose and spun again at 10,000x g for 15 min to yield the washed synaptosomal fraction. The resulting pellet was lysed by hypoosmotic shock in 9 volumes ice cold H_2_O plus protease/phosphatase inhibitors and the HEPES concentration was rapidly adjusted to 4 mM and further incubated at 4°C for 30 min to ensure complete lysis. The lysate was centrifuged at 25,000x g for 20 min to yield the crude synaptic vesicle fraction and the synaptosomal plasma membrane. The pellet was resuspended in HEPES-buffered sucrose and layered on top of a discontinuous gradient containing 0.8 to 1.0 to 1.2 M sucrose supplemented with protease/phosphatase inhibitors. The gradient was centrifuged in a swinging bucket rotor (SW41) at 30,000 rpm for 2 hr. The synaptic plasma membrane was recovered in the layer between 1.0 M and 1.2 M sucrose.

### Determination of the F/G-actin ratio

The F/G actin ratio determination was performed as described earlier (Lopes et al., 1999; Rust et al., 2010). Freshly dissected brain tissue from P30 mice was lysed in ice cold 1xPHEM extraction buffer (60 mM PIPES, 20 mM HEPES, 10 mM EGTA, 2 mM MgCl_2_ and 1% Triton X-100; pH 7.0) by using a tight fitting Dounce homogenizer. F-actin was pelleted by a 10 min centrifugation at 10,000 rpm. Supernatant and pellet were brought to equal volume by 1x SDS sample buffer, heated at 95°C and equal amounts of each sample were separated on 10% polyacrylamide SDS gels, transferred to nitrocellulose membranes, and subjected to immunolabeling.

For F/G actin analysis, synaptosomes from forebrain of P30 mice were isolated as previously described (Villasana et al., 2006). Forebrain tissues were homogenized in synaptosome extraction buffer (10 mM HEPES, 1 mM EDTA, 2 mM EGTA, 0.5 mM DTT, 10 µg/ml Leupeptin, 20 µg/ml Aprotinin; pH 7.0) using a Teflon glass mechanical tissue grinder on ice. The homogenate was sonicated and loaded into a 60 ml Luer Lok syringe and filtered twice through a 100 µm pore nylon filter (Millipore). The resulting filtrate was filtered through a prewetted 5 µm pore hydrophilic filter (Millipore) through a 5 ml Luer Lok syringe and centrifuged at 10,000x g and 4°C for 10 min. Synaptosomes were resuspended and lysed in ice cold 1x PHEM extraction buffer by using a tight fitting douncer. F-actin was pelleted by a 10 min centrifugation at 10,000 rpm. Supernatant and pellet were brought to an equal volume by 1x SDS sample buffer, heated at 95°C and equal amounts of each sample were separated on 10% polyacrylamide SDS gels, transferred to nitrocellulose membranes, and then subjected to immunolabeling.

For KCl depolarization assay, synaptosomes were resuspended in Krebs buffer, equilibrated and incubated with 40 mM KCl (Sigma-Aldrich) for the indicated time points. F and G actin was fractioned using PHEM extraction buffer as described above. Anti-β actin was used as primary antibody (1:2500; Sigma), and HRP-conjugated anti mouse secondary antibody (1:10000) was used for detection. Densitometric analysis was done using Image J for estimation of the F/G-actin ratio.

### Western blot analysis

Brain tissues from embryonic (E18.5), juvenile mice (P30) and adult mice (P365) were homogenized and lysed (1% Triton X-100, 0.1 M NaCl, 0.05 M Tris-HCl, pH 7.5, 10 mM EDTA, freshly added 1x protease inhibitor cocktail (PIC), 0.5 mM PMSF, 0.01 M DTT).

Dissected cortices from E18.5 embryonic brains were resuspended by pipetting in cold extraction buffer (1% Triton X-100, 0.5% Nonidet P-40, 25 mM Tris-HCl, pH 7.5, 100 mM NaCl) supplemented with 1X EDTA free protease inhibitor cocktail (Roche, Mannheim, Germany), 50 µg/ml 4-(2-aminoetyl)benzensulfonyl fluoride hydrochloride (Sigma) and 1 mM sodium orthovanadate (Sigma). After mixing for 30 min at 4°C, the samples were centrifuged for 10 min at 16,100 x g, and the supernatant transferred to a fresh tube. For the LIM kinase and phospho-LIM kinase blots lysates were prepared by resuspending dissected cortices in 20 µl/mg wet tissue weight of the following buffer: 10 mM Tris, pH 7.5, 1% SDS, 0.5 mM DTT, 2 mM EGTA, 20 mM NaF, 1 mM Na_3_VO_4_ and protease inhibitors. After sonication for 3 s, the lysates were plunged into a boiling water bath for 10 min (Garvalov et al., 2007).

SDS-sample buffer was added to the extracts, followed by heating at 95°C for 5 min. Proteins were resolved on 10% and 16% polyacrylamide SDS gels, transferred to nitrocellulose membranes, and then subjected to immunolabeling. Primary antibodies used were polyclonal anti-CAP2 (1:800), polyclonal anti-CAP1 (1:1200) (Peche et al., 2007), monoclonal anti-β actin (1:2500; Sigma), monoclonal anti-cofilin (1:1000; Cell signaling), monoclonal anti-phospho cofilin (1:750; Cell signaling), monoclonal anti-LIMK (1:750; Cell signaling), monoclonal anti-phospho LIMK (1:750; Cell signaling), monoclonal anti-TESK1 (1:1000; Cell signaling), monoclonal anti ROCK1 (1:1000; Cell signaling), monoclonal antichronophin (1:1000; Cell signaling). HRP-conjugated anti mouse or anti rabbit secondary antibodies (1:10,000) were used for detection. Monoclonal antibodies against GAPDH conjugated with horseradish peroxidase (Sigma, St. Louis, MO, USA) were used for analyzing equal loading. Densitometry analysis was done using Image J for estimation of the phospho cofilin.

### **Golgi staining** (Glaser and Van Der Loos, 1981)

Mice at P30 were used for Golgi-Cox staining in order to study the dendritic spine density using the FD Rapid Golgi Stain kit (FD Neurotechnologies). Tissue impregnation and tissue section staining were performed according to the manufacturer’s protocol. Briefly, mice were perfused with 4% paraformaldehyde (PFA) and post fixed in 4% PFA overnight. After incubation in impregnation solution and solution C, brains were cut into 150 µM coronal sections using a Vibratome (Leica, Germany). Sections were mounted onto gelatinized glass slides and further processed for Golgi staining procedure and finally mounted in permount (Sigma). High magnification images of dendritic branches from the cortex and hippocampus were generated by using an Axioskop microscope and a Plan-Neofluar x100/1.30 oil immersion objective (Carl Zeiss). The images were then reassembled and the dendritic spine analysis was performed using RECONSTRUCT software as described previously (Risher et al. 2014).

### Analysis of Purkinje cell morphology

The animals were anesthetized with halothane (B4388; Sigma-Aldrich, Taufkirchen, Germany) and subsequently decapitated. Coronal slices (300 μm) were cut with a vibration microtome (HM-650 V; Thermo Scientific, Walldorf, Germany) under cold (4 °C), carbogenated (95% O2 and 5% CO2), glycerol-based modified artificial cerebrospinal fluid (GaCSF) (Ye et al., 2006) to enhance the viability of neurons. GaCSF contained 250 mM Glycerol, 2.5 mM KCl, 2 mM MgCl_2_, 2 mM CaCl_2_, 1.2 mM NaH_2_PO_4_, 10 mM HEPES, 21 mM NaHCO_3_, 5 mM Glucose and was adjusted to pH 7.2 with NaOH resulting in an osmolarity of 310 mOsm. Brain slices were transferred into carbogenated artificial cerebrospinal fluid (aCSF). First, they were kept for 20 min. in a 35 °C ‘recovery bath’ and then stored at room temperature (24 °C) for at least 30 min prior to recording. Neurons were visualized with a fixed stage upright microscope (BX51WI, Olympus, Hamburg, Germany) using 40× and 60× water-immersion objectives (LUMplan FL/N 40×, 0.8 numerical aperture, 2 mm working distance; Olympus) with infrared differential interference contrast optics and fluorescence optics (Dodt and Zieglgansberger, 1990). 1% Biocytin and tetramethylrhodamine-dextran was allowed to diffuse into the single cell until dendritic arborizations of Purkinje cells were clearly visible. The brain slices were fixed in Roti-Histofix (P0873; Carl Roth, Karlsruhe, Germany) overnight at 4°C and rinsed in 0.1 M PBS afterwards and subsequently incubated in Alexa Fluor 633 (Alexa 633)-conjugated streptavidin (1:600; 2 hours; RT; S21375; Invitrogen, Karlsruhe, Germany) that was dissolved in PBS containing 10% normal goat serum. Images were obtained using a Zeiss LSM 510 confocal laser scanning microscope with an Achroplan 40x/0.75 oil immersion objective (Zeiss). A stack of optical sections (1 µm) completely covering the labeled cell was acquired. The stacks were reassembled and the morphometric analyses were performed with ImageJ. A concentric Sholl analysis and reconstruction of neuronal structure was performed with NeuronStudio (Beta) as reported earlier (Rodriguez et al., 2008; Wearne et al., 2005). At least four animals were used for each data set.

### Cofilin depletion assay

The supernatant depletion pull down assay for examining the interaction between GST-tagged CAP2 and thrombin released cofilin from GST-cofilin was carried out as described (Mattila et al., 2004). GST-CAP2 fusion proteins were expressed in *E. coli (BL21)* and harvested. The resulting pellet was resuspended in lysis buffer (10 mM Tris, pH7.5, 100 mM NaCl, 1 mM DTT, 4 mM EDTA, 0.1% Triton X-100, 0.2 mM PMSF), lysed by sonication and centrifuged at 18,000 x g for 30 min at 4°C. The fusion protein was immobilized on glutathione-sepharose beads (Macherey-Nagel) at a concentration of 0.1, 0.2, 0.5, 1 and 2 µM. GST was removed by thrombin from GST-cofilin. Equal volumes of the cofilin solution were incubated with GST or GST-CAP2 coupled to glutathione-agarose beads, and the amount of cofilin in the supernatant was quantified from Coomassie-Blue stained SDS-polyacrylamide gels by densitometry using Image J.

### Pull-down assays

GST-CAP2_FL_, GST-CAP_256-476_, GST-N-CAP2WH2, GST-WH2, GST-WH2-C-CAP2 and GST-ΔWH2-CCAP2 cloned in pGEX 4T-3 were expressed in *E. coli (BL21)*. The fusion proteins were immobilized on glutathione-sepharose beads. GFP-cofilin was transiently expressed in HEK293T cells. For transfection of the corresponding plasmids Lonza transfection kit was used. Cells were harvested next day, lysed in lysis buffer and equal amounts cell lysates were incubated with GST fused domains of CAP2 for 2 hrs at 4°C. Precipitated proteins were detected by western blotting using mAb K3-184-2 specific for GFP (Blau-Wasser et al., 2009).

For pull down of transiently expressed GFP-CAP2 from HEK293 cells, the cell lysate was incubated with GST-cofilin WT, GST-cofilin S3A and GST-cofilin S3D for 2 hrs. at 4°C. The beads were washed and resolved on 10% polyacrylamide SDS gels and pull down probed with mAb K3-184.2.

### Statistical Analysis

Set of data are presented by their mean values and standard error of the means. An unpaired two tailed Student’s *t*-test was used when comparing 2 sets of data using Microsoft excel. Data are presented as mean ±standard error of the mean.

## Acknowledgements

VSP acknowledges support by “Köln Fortune” and the “Maria Pesch Foundation”. AK acknowledges support from “Imhoff-Stiftung”. Authors thank the CECAD imaging facility, Cologne, and DZNE imaging facility, Bonn. We thank Dr. Walter Witke for n-cofilin construct. There are no financial disclosures or conflicts of interest existing for any of the authors.

## Author contributions

AK, performed experiments, design, data analysis and interpretation, manuscript writing; LP, Purkinje neuron labeling, KK, helped in immunofluorescence analysis; PK, experimental design; AAN, conception and design, manuscript writing and fundraising; VSP, conception and design, data analysis and interpretation, manuscript writing and fundraising. All authors reviewed the manuscript. The authors declare no competing financial interests.

---

**Supplementary Figure 1:**
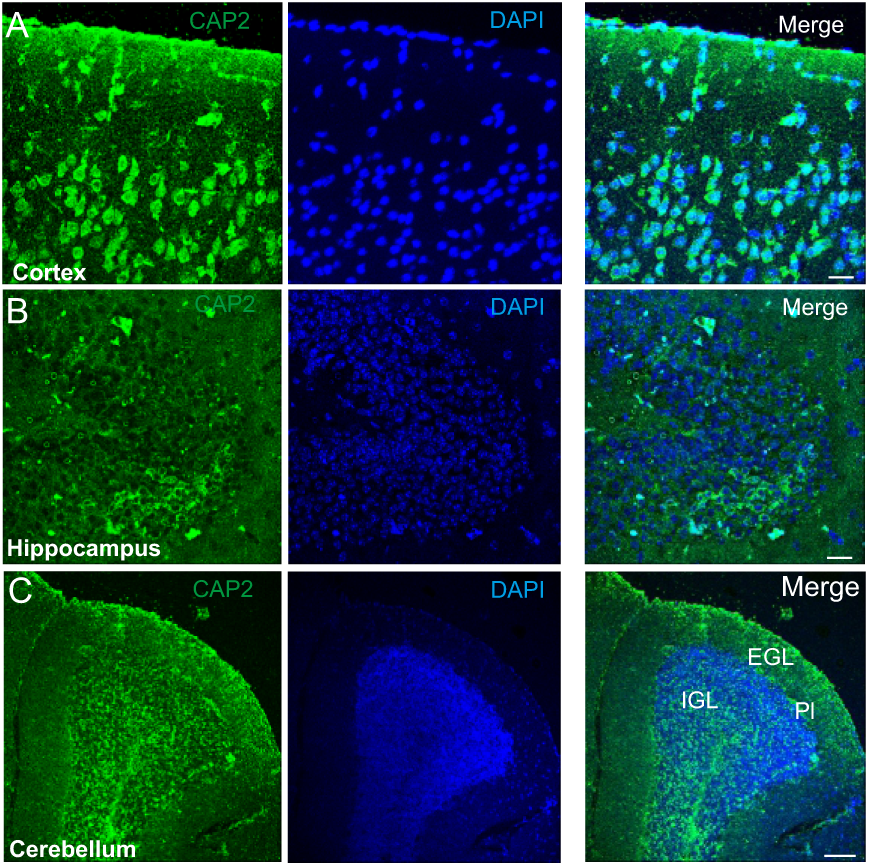
**(A,B,C)** CAP2 is expressed in cortex, hippocampus and cerebellum. Paraffin embedded brain section was stained with CAP2 monoclonal antibody and widespread expression profile was observed in mice brain. (Scale bar: 20µm).

**Supplementary Figure 2:**
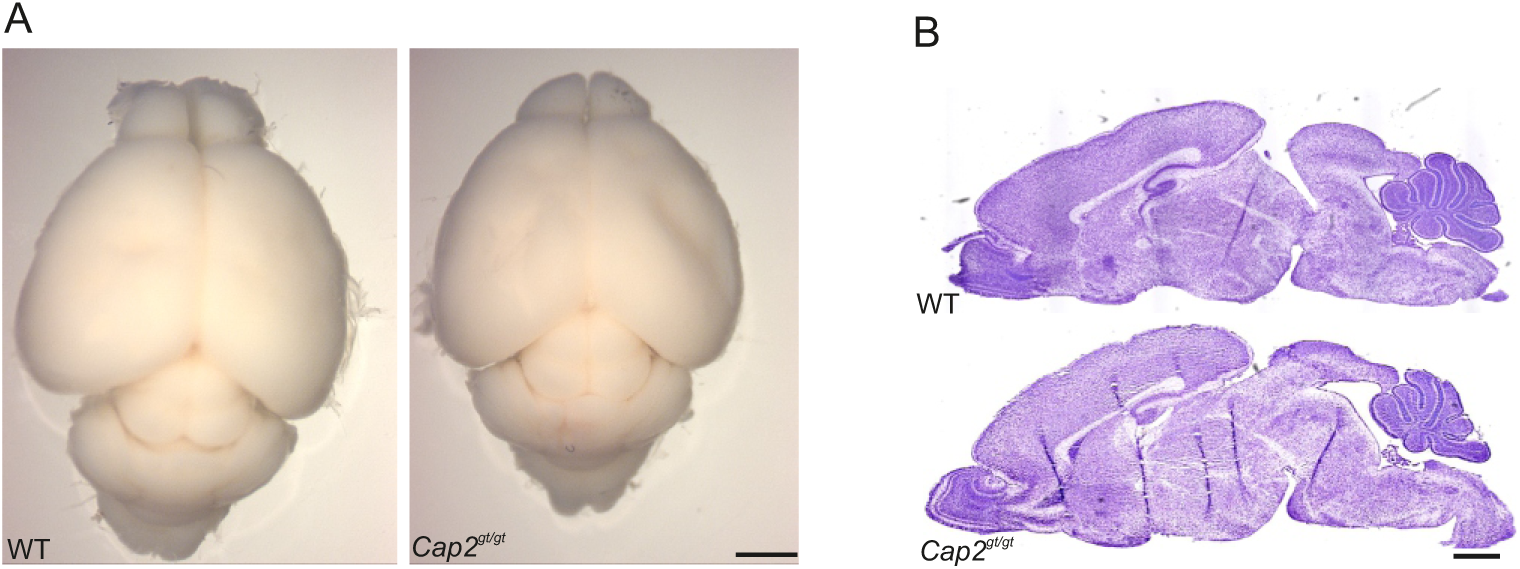
**(A)** No significant difference in gross anatomy of brain at post natal day 8 was observed (Scale bar: 1mm) (**B)** Cresyl violet staining does not reveal any change in organization of brain. Sagittal sections from control and mutant brains are depicted (Scale bar: 250µm).

**Supplementary Figure 3:**
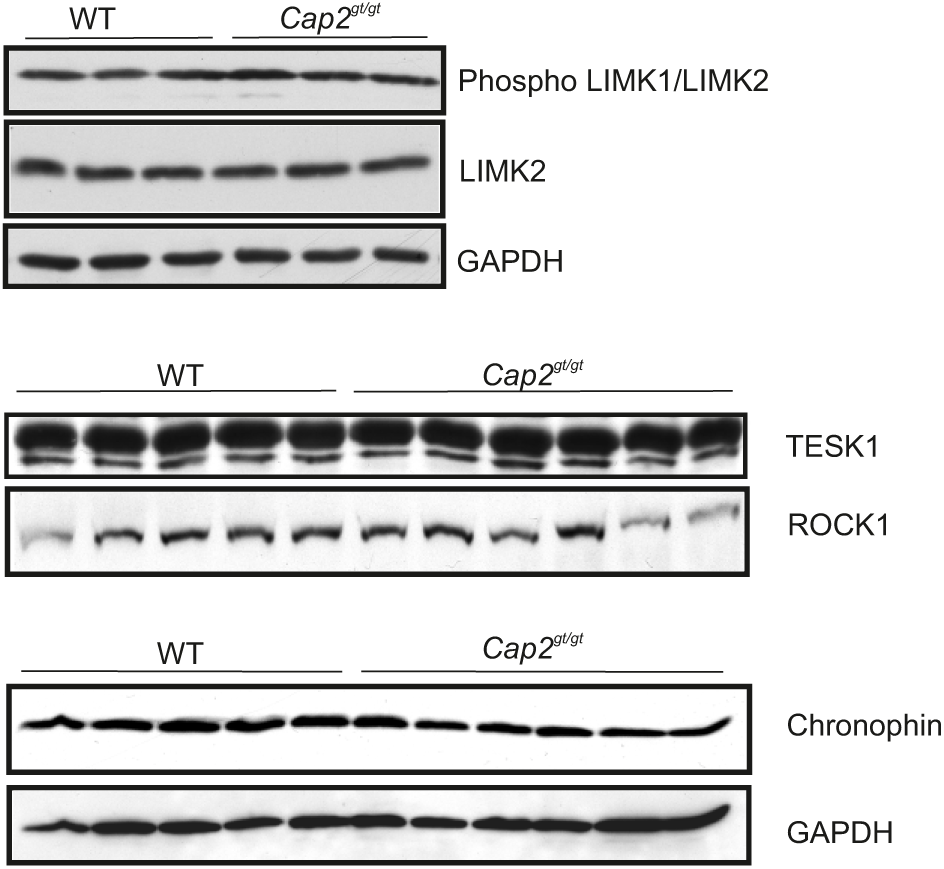
Expression of kinases and phosphatases involved in Cofilin phosphorylation/dephosphorylation cycle. Western blotting of phospho LIMK1/LIMK2 and LIMK2 do not show any significant difference in WT and CAP2 mutant brain lysate. The levels of other kinases like TESK1 and ROCK1 were also unaltered in mutant brain lysate as was the level of chronophin.

**Supplementary Figure 4:**
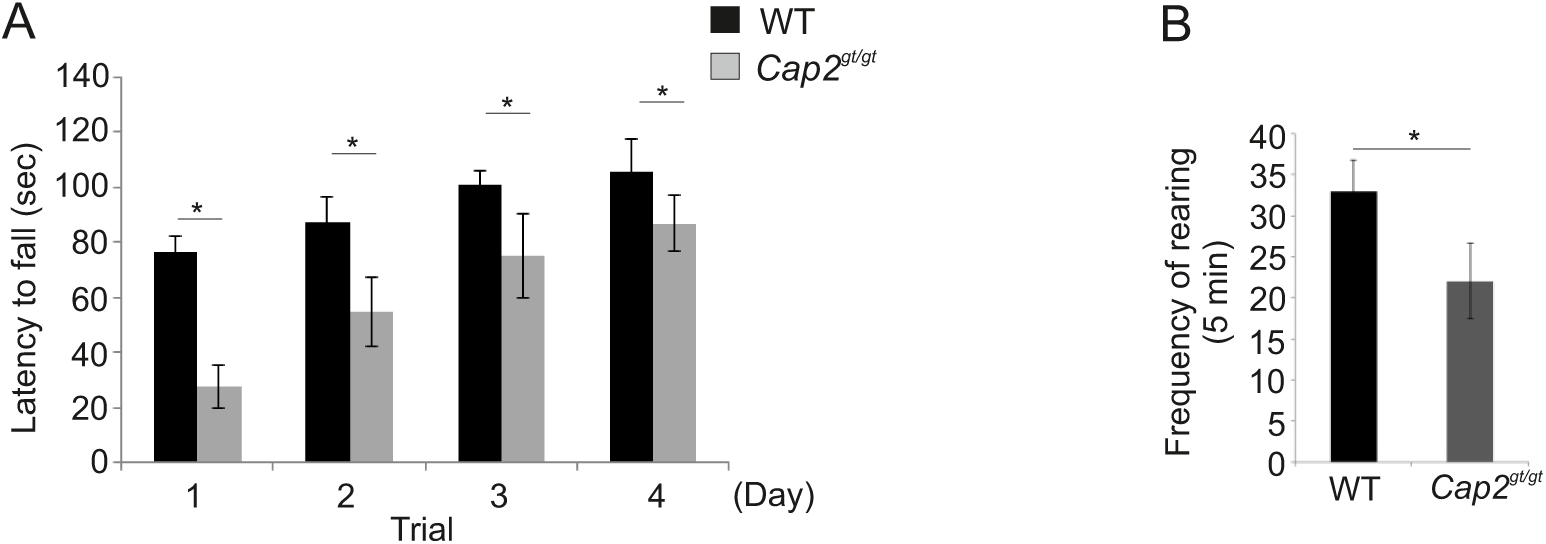
**(A)** The latency to fall from an accelerating rota rod from 4 to 40 seconds was measured from three trials per day (Trial 1: WT = 76.45± 5.8; n= 8; *Cap2^gt/gt^* = 27.72 ± 7.8; n=12; P= 0.001; Trial 4: WT = 105±12; n= 8; *Cap2^gt/gt^* = 86± 10; n=12; P= 0.02). **(B)** Rearing frequency in an open field test was observed in WT and Cap2 mutant mice (WT = 33± 3.2; n= 8; *Cap2^gt/gt^* = 21± 4.6; n=12; P= 0.036).

